# Assessing the degradation dynamics of sugar kelp in anaerobic marine sediment using environmental DNA

**DOI:** 10.64898/2026.06.23.734019

**Authors:** Samuel H. Tan, Robin S. Sleith, Nichole N. Price, David Emerson, Jeremy J. Rich

## Abstract

Environmental DNA (eDNA) has the potential to be a powerful tool in blue carbon science for characterizing and quantifying the contribution of marine macrophytes; but its complex, dynamic relationship with bulk biomass is poorly understood. Here, we used eDNA to examine the degradation dynamics of sugar kelp (*Saccharina latissima*) in muddy, anaerobic marine sediment. This involved three 16-week incubations; with additions of lyophilized sugar kelp alone, a mix of lyophilized marine macrophytes including sugar kelp, and sugar kelp holdfasts buried in sediment. We used species-specific digital polymerase chain reaction assays for mitochondrial, chloroplast and nuclear markers, and metabarcoding for the 16S and 18S ribosomal RNA genes. In the former two incubations, all sugar kelp eDNA markers showed rapid log exponential declines (up to 98-99%) to asymptotes greater than the unamended controls, even as part of a more complex mix of macrophytes. In contrast, for the buried kelp holdfasts, sugar kelp eDNA increased to an asymptote (by up to ∼15X), which may be reflective of the different nature of added biomass. Overall, we demonstrate substantial preservation of environmental DNA and total organic carbon under anaerobic conditions, and the potential to use environmental DNA to quantify biomass in a blue carbon context.

## Introduction

A central challenge for understanding the degradation of macroalgal biomass entering marine sediments is being able to track its fate within the complex sediment matrix ^1–3^. This is relevant both to understanding the fate of natural seaweeds in coastal sediments ^4,5^, and as a method for carbon accounting related to blue carbon proposals to grow kelp and other macroalgae and then store their carbon in either sediments or the deep oceans as a means of marine carbon dioxide removal (mCDR) ^6–8^. Environmental DNA (eDNA) has the potential to be a powerful tool in blue carbon science for estimating the contribution of macroalgae (and other macrophytes) to organic matter (OM) in marine sediments ^9–12^, with its strengths of high sensitivity and specificity at potentially lower costs than most biogeochemical methods ^9,13^.

eDNA methods have detected marine macroalgal biomass in water and sediment samples ranging from coastal waters to the deep sea ^11,14–16^, including our recent work ^17^, and demonstrated some degree of connectivity between potential source (e.g. macroalgal beds) and sink sites (e.g. soft sediments). However, mCDR strategies, including the promising strategy of sinking aquacultured marine macroalgal biomass ^7,18^, require Measurement, Monitoring, Reporting and Verification (MMRV) ^2,18^ to reliably track and quantify stored carbon. One of the key knowledge gaps limiting eDNA from quantitative use as a powerful component of MMRV toolkits is a lack of knowledge about the complex, often context-dependent degradation dynamics of eDNA in the environment ^13,19,20^, which is specific to both the form of eDNA (e.g. tissues, cells, free and complexed DNA) and study system (e.g. air, sediment, water [freshwater & marine]) Hence, the actual quantity of biomass which a unit of eDNA (e.g. gene copies, read counts) represents, and how that relationship may change over time as biomass degrades and ages is not well understood ^13,20^. Even less is known about the degradation dynamics of eDNA in marine sediments ^21^, especially fine, anaerobic sediments where even labile OM can be well preserved ^22,23^.

To date, the majority of studies have focused on the effect of macroalgal addition on sediment nutrient cycling ^24,25^ and the associated microbiome ^26–28;^ but rarely aim to characterize the fate of macroalgal biomass per se ^29–32^, with some exceptions ^16,25,33^. These studies tend to rely on biogeochemical markers (e.g. stable isotopes, fatty acids ^16,25^) which are less species-specific than eDNA ^9,10^. A substantial amount of{Citation} sinking macroalgal biomass is likely to make it to the seafloor ^4,29,34,35^. In deeper waters, even if this biomass is remineralized, the resultant inorganic carbon can remain in deep ocean circulation for climate-relevant timeframes (i.e. hundreds of years) ^34,36^. Depending on location, a large proportion of sinking biomass may make it to shallower sediments ^16^, which are much more prone to disturbance from biological, physical and anthropogenic sources, but may still be able to accumulate significant amounts of organic carbon if deposition and burial rates exceed remineralization ^37^.

Thus, our aim in this study is to examine how sugar kelp (*Saccharina latissima*) eDNA from added biomass changes over time in anaerobic sediments, and how multiple eDNA markers from different genomic and/or organellar sources, both quantitatively (i.e. dPCR) and qualitatively (i.e. metabarcoding) correlate with one another and bulk total organic carbon (TOC) and total nitrogen (TN) over time. Additionally, we tested the hypothesis that comparing the recovery ratios of longer to shorter gene fragments and targets of differential starting abundance can provide a proxy of biomass age ^38–41^. We compared the degradation dynamics between different types of added *S. latissima* biomass, with finely ground biomass; and kelp holdfasts representing non-commercially viable material that could be sunk for mCDR purposes ^18^. Overall, we hope to use eDNA to shed light on the early degradation dynamics and fate of macroalgal biomass in marine sediments.

## Results

### Relationship between added biomass and gene copy numbers for the 4 assays

The four dPCR assays of different eDNA targets (150bp & 300bp COI, 100bp rbcL, and 120bp NC fragments) performed well, showing strong linear relationships between gene copy number and added *S. latissima* biomass (Table S2 & Figure S1). In general, the rbcL assay was the most sensitive (p < 0.0001), with the highest gene copies per unit of added biomass (about twice as sensitive as the 150bp COI assay). The 150bp COI assay was about 1.5x as sensitive as the 300bp COI assay (p < 0.0001), and 60x more sensitive than the NC assay (p < 0.0001). The results suggest that these gene targets serve as suitable proxies for *S. latissima* biomass in the degradation experiments, more so for rbcL and 150bp COI.

### LBA: kelp experiment: Degradation dynamics of eDNA and non-eDNA markers

In 16-week incubations of added lyophilized sugar kelp alone (LBA:kelp experiment), all samples were analyzed using the four dPCR assays mentioned above, and two semi-quantitative eDNA assays: 16S read counts for chloroplasts (of which the vast majority in the kelp treatment [>95%] were verified as originating from *S. latissima*, using BLAST), and 18S read counts for *S. latissima*; as well as the non-eDNA metrics, added TOC% and added TN% (Figure 1). All of the standalone eDNA metrics (excluding the 150bp:300bp COI ratio), showed log exponential declines that then levelled off after about 6-8 weeks and remained at stabilized levels that were significantly higher than the controls (e.g. ∼62X for 150bp COI, ∼17X for 300bp COI, ∼139X for rbcL, ∼5X for NC, comparing Week 16 values with mean values for the controls), for the rest of the incubation (Figure 1, Figure S2, Tables S3-S4). The mitochondrial (COI) and chloroplast dPCR markers (rbcL) declined by ∼98% on average from the start of the incubation to stabilized levels that were ∼2% of initial dPCR gene copies (Table S4) after 6-8 weeks. The dPCR eDNA indicators, 150bp COI, 300bp COI and rbcL correlated strongly with one another throughout the experiment (Table S5). 18S *S. latissima* read counts and 16S chloroplast read counts also showed significant correlations with the above dPCR eDNA indicators and decreased by ∼87% and ∼99% from initial values respectively. The 150bp:300bp COI ratio (Figure 1, Figure S3, Table S6) increased over time to an asymptote (by about ∼1.5X). The NC assay showed weaker correlations with the other assays, as well as 18S and 16S, likely due to the fact that there are only two copies of the NC marker per kelp cell (in a diploid adult sporophyte), while there are many copies of mitochondrial (COI) and chloroplast (rbcL) genes, as well as ribosomal RNA genes (18S in the nucleus, and 16S in the chloroplasts) per cell.

**Figure 1:**
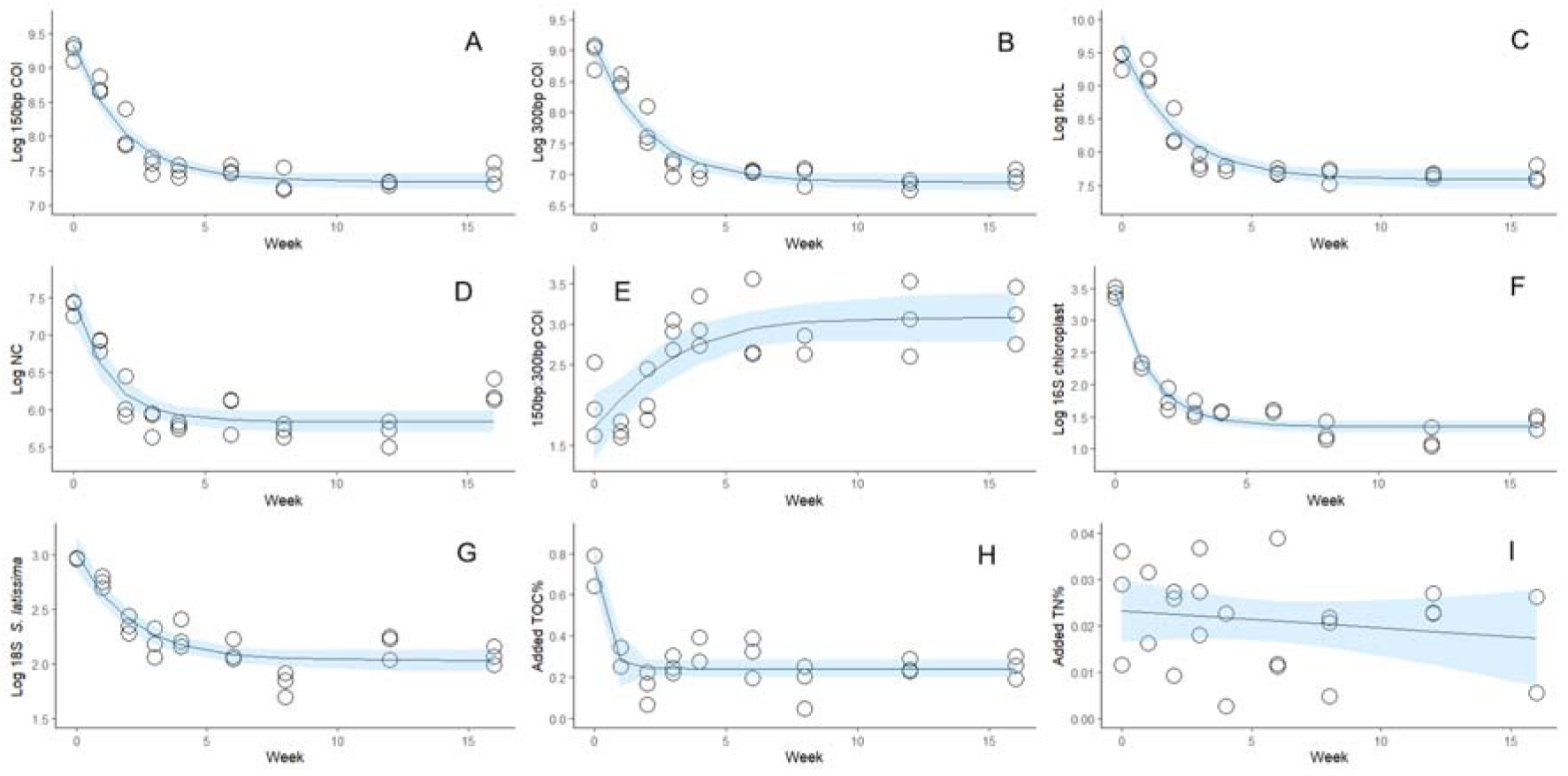
Changes over time in the LBA: kelp experiment in four dPCR markers of *S. latissima* (A-D), the ratio between two of these markers (E); the two metabarcoding markers (F & G; log 16S chloroplast read counts and log 18S *S. latissima* read counts respectively); and added TOC% (H) and added TN% (I). The line in each graph corresponds to average model fits with 95% confidence intervals in light blue. Note that y-axis values are not all the same, and that the eDNA markers (A-D, F-G) use a log scale for a y-axis.

Given that the only added biomass in the LBA: kelp experiment was *S. latissima*, the markedly greater 16S chloroplast reads compared to the controls were assumed to correspond primarily to this taxon, even though we did not expect this marker to have the taxonomic resolution to distinguish chloroplasts from different taxa. Added TOC% levels (i.e. after subtracting mean values in the controls) also showed an initial rapid decline and were always significantly greater than zero. However, in contrast to the larger overall decreases in COI and rbcL gene copies over 6-8 weeks, added TOC% decreased and stabilized at about 35% of initial levels by 2-4 weeks. Added TN% values were also significantly greater than zero throughout the experiment (Figure 1, Figure S2, Table S3).

### LBA: mixed experiment: eDNA metrics and degradation dynamics of macrophyte 18S reads

Unlike in the LBA: kelp experiment, here we examined the degradation dynamics of *S. latissima* eDNA in the presence of a uniform mixture of 7 other different macrophyte taxa from a wide range of taxonomic groups (for a total of 8 taxa). In this experiment, we used 18S read counts to assess overall changes in relative abundance of the different taxa, and the dPCR 150bp COI assay to specifically track the fate of *S. latissima*.

Similar to the LBA: kelp experiment, dPCR gene copies of 150bp COI showed log exponential declines to an asymptote with end values significantly greater than the controls. Given the smaller proportion of added *S. latissima*, starting and ending values for 150bp COI gene copies (∼1.7X lower initially, ∼37X lower at the end) and 18S *S. latissima* reads (∼3X lower initially, ∼12X at the end) were lower than in the LBA: kelp experiment. In contrast, the total number of 16S chloroplast reads was higher in this experiment. This result is best explained by the fact that the total number of chloroplasts originating from the multiple added macrophyte taxa was substantially greater than in the kelp-only experiment. (Figure S2, Table S3). While 18S degradation dynamics for *S. latissima* were similar to the LBA:kelp experiment, there were notable differences in how 18S gene copies changed in other macrophyte taxa. Four taxa (F*. vesiculosus, A. nodosum, C. crispus* & *Rhodomela* sp.) showed significant log linear declines (Figure 2 & Table S7), while linear model fits for *Ulva* sp. & Z. *marina* were non-significant. Interestingly, *Z. marina*, the only embryophyte, showed greater mean read counts at the end of the experiment, which suggests that it was relatively more resistant to degradation than the other taxa.

**Figure 2:**
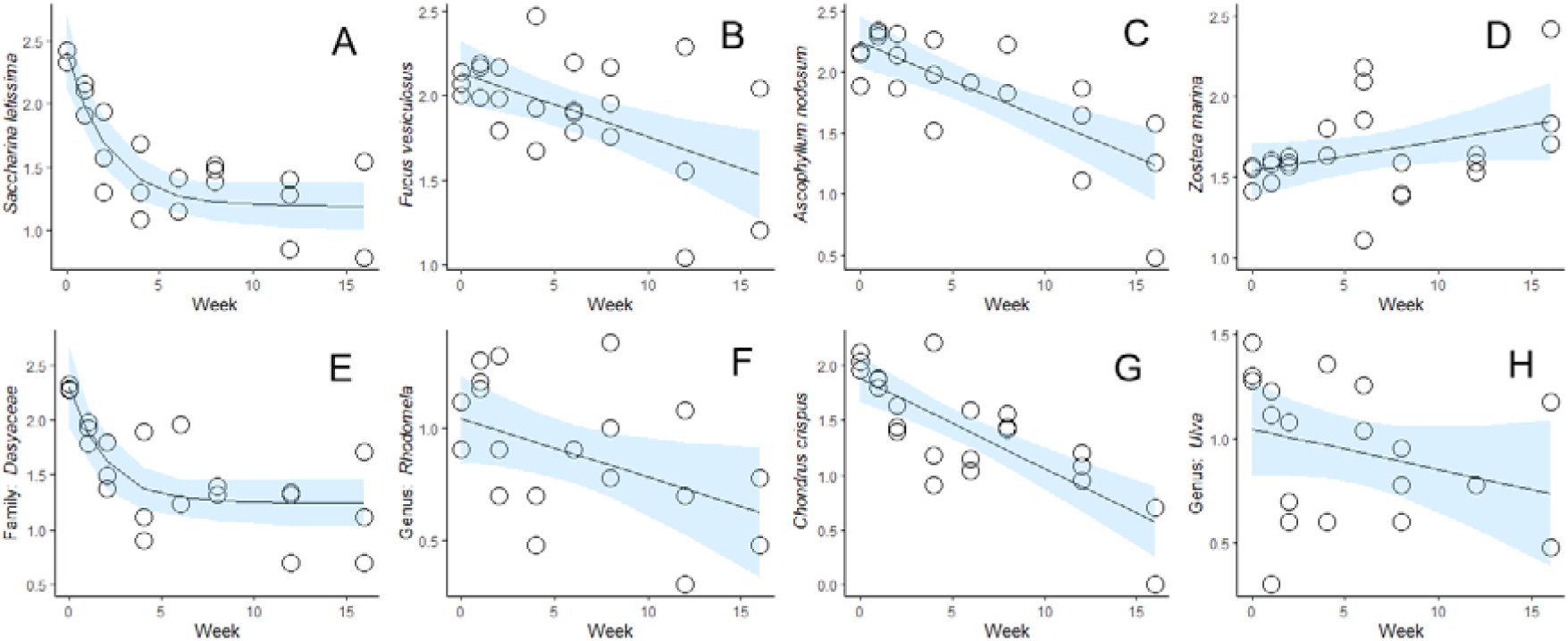
Changes in log 18S read counts over time for the 8 added macrophyte taxa in the LBA: mixed experiment. The line in each graph corresponds to average model fits with 95% confidence intervals in light blue. Note the y-axis values are not all the same.

### KB experiment

In contrast to the above experiments, we examined the change in eDNA released from intact *S. latissima* holdfasts buried in previously undisturbed sediment (Image S1) by measuring eDNA in the surrounding bulk sediment. Interestingly, numbers of both *S. latissima* COI gene copies (∼15X) and 18S read counts (∼3.8X) increased during the first four weeks of the experiment, and then plateaued (Figure 3, Figures S4-S5, Table S9). This is a marked contrast to the decline in these markers in the LBA experiments. Correspondingly, the bulk TOC% and TN% of surrounding sediments were significantly higher in holdfast treatments compared to controls, although the differences were more variable and smaller than in the LBA experiment (Figures S4 & Table S8). During the experiment, the holdfasts became limp and lost approximately 80% of their mass (Figure S5F), although they retained basic structural integrity.

**Figure 3:**
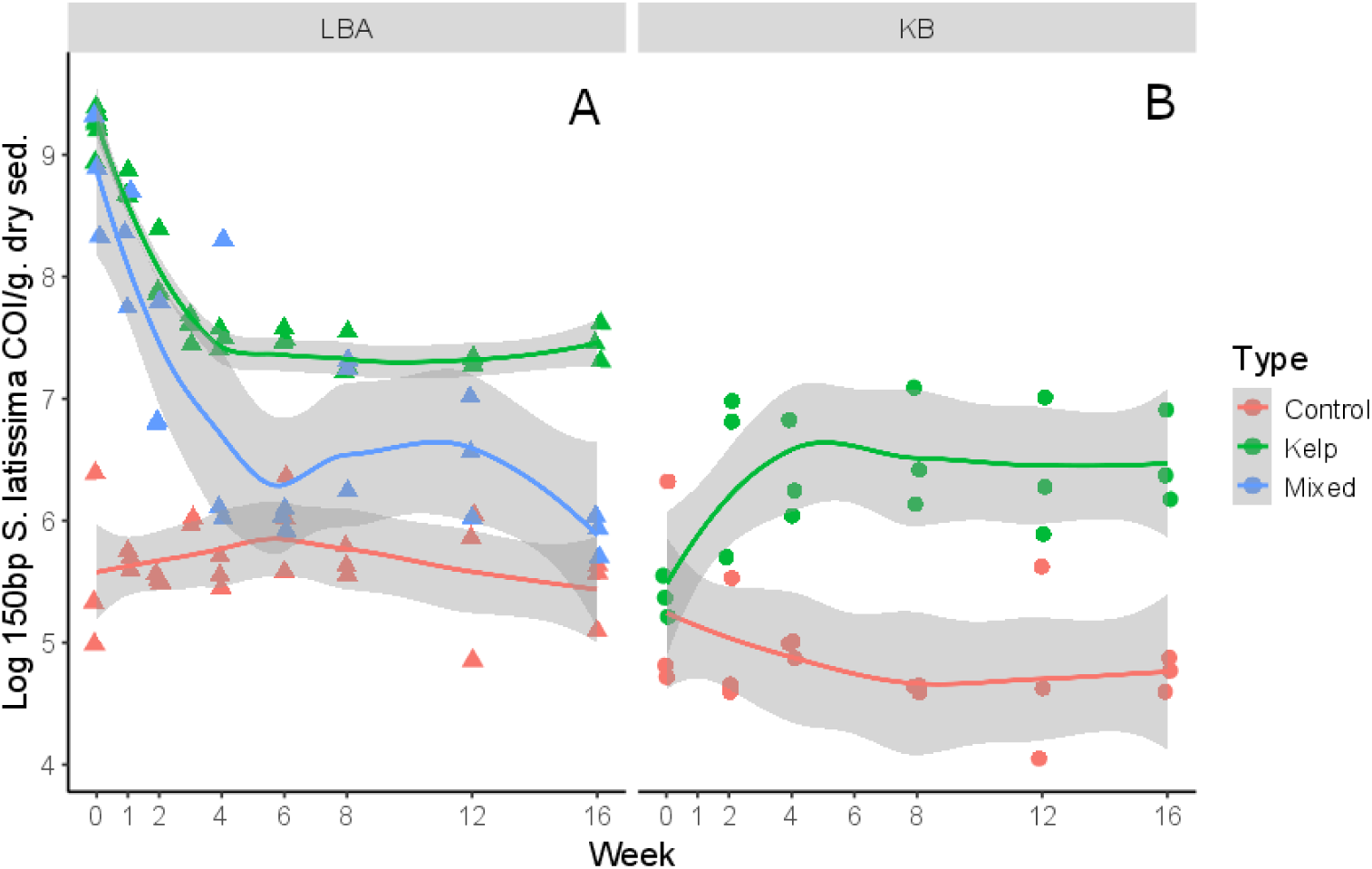
Log 150bp *S. latissima* COI gene copies in the KB experiment (Panel B) (results from both LBA experiments combined [Panel A] are shown again for a direct comparison). Smoothed means have been drawn through the points for each treatment type for each metric, with the surrounding gray outline representing the 95% confidence interval.

## Discussion

Our study shows that in undisturbed anaerobic sediments, burial of kelp and macrophyte biomass can result in substantial preservation of OM. This supports the idea that buried macrophyte biomass can contribute to carbon sequestration in marine sediments under appropriate conditions. Furthermore, our study reinforces the idea that eDNA can be a highly sensitive, specific and low-cost tool for tracing and characterizing the fate of macrophyte biomass in marine systems ^9–11^. In a blue carbon context, eDNA can be an important component of a complementary suite of biogeochemical and molecular biological tools required to reliably trace macrophyte OM across complex seascapes

To establish a basis for using eDNA to estimate *S. latissima* degradation rates, we demonstrated strong linear relationships between added *S. latissima* biomass and gene copy number for all four dPCR assays in non-incubated, spiked sediments. This is consistent with our previous study ^17^, but adds three additional gene markers (rbcL, 300bp COI, and NC) that may be used. Of these additional markers, rbcL was the most sensitive, consistently yielding the greatest number of gene copies per unit of *S. latissima* biomass. This work also shows that metabarcoding read counts using the 18S gene correlate well with gene copies from quantitative dPCR assays, although metabarcoding is less sensitive than dPCR ^42^. This relationship also becomes more complex as biomass degrades.

Our LBA experiments show that based on the dPCR data, ∼98% of the *S. latissima* eDNA that was initially present was rapidly degraded in a log exponential fashion within six weeks of the experiment. It is generally known that labile OM in marine sediments can degrade at similar rates under aerobic and anaerobic conditions in marine sediments ^43–46^. However, few studies have examined the degradation of macroalgae in anaerobic marine sediments ^33^, and unlike this study, solely rely on biogeochemical measurements to estimate decay rates. In this study, most eDNA derived from freshly added kelp material would likely represent among the more labile fractions of OM as DNA can be readily hydrolyzed if unprotected from enzymatic degradation ^47^. In kelp, other labile OM species may include alginate, laminarin, mannitol; and various proteins and amino acids. Over time, the remaining OM pool is expected to be increasingly represented by more recalcitrant species and thus becomes more difficult to degrade under anaerobic conditions compared to aerobic conditions ^48,49^. In kelp, these recalcitrant OM species may include complex structural polymers that are difficult to hydrolyze without complex suites of enzymes ^50^, such as sulphated polysaccharides (e.g. fucoidans) and polyphenolics (e.g. phlorotannins) ^32^. Despite the high eDNA degradation rates in our study (up to ∼100-fold over the course of 16 weeks, with eDNA T_50_ values [i.e. time to taken to fall to 50% of the initial eDNA gene copies] for all dPCR targets being on the order of days [i.e. comparable to aerobic aquatic and marine settings ^39,40^]; Table S4); most of these eDNA metrics eventually levelled off for the duration of the experiment and remained higher than controls. This remaining eDNA (and more broadly, labile OM) may be preserved through mechanisms such as complexation with minerals (e.g. clays) and more recalcitrant forms of complex OM (e.g. humics), which are known to allow labile OM to persist in substantial amounts in marine sediments over long time periods ^21,22,51^. This phase of greatly reduced degradation is also seen in the ratio between 150bp and 300bp COI fragments. Longer eDNA fragments are expected to degrade faster than shorter fragments as they are less likely to be protected by complexation with minerals and/or organics, and present larger targets for hydrolysis ^20^. The stabilization of this ratio by Week 4-6 at around ∼3X (Figure S3) indicates that degradation has slowed to the extent where innate differences in degradability have little effect. Overall, this supports the paradigm that substantial preservation of even labile OM can still occur under anaerobic conditions ^52,53^, including DNA ^23^.

Given the relatively short 16-week timeframe of our experiment, even longer-term studies may be needed to examine eDNA persistence in this plateau phase of lower degradation rates ^13,54^. We caution that these findings were determined under stable laboratory conditions, while in natural sediments, eDNA may be exposed to varying oxic-anoxic conditions due to processes like bioturbation or sediment resuspension, which would further promote degradation. Nevertheless, given that a readily detectable eDNA signal persisted along with significant preservation of bulk added TOC in the sediments, this study suggests that eDNA techniques could be used to detect *S. latissima* biomass, even after substantial degradation of labile OM.

We also looked at eDNA degradation in a more diverse mix of marine macrophytes, but general eDNA degradation dynamics of *S. latissima* remained similar to the kelp only treatment. This indicated that the presence of OM from multiple macrophyte taxa does not have synergistic or antagonistic effects on the degradation of *S. latissima*. With the exception of log exponential declines in 18S read counts of *S. latissima* and *Dasyaceae*, all other taxa (except *Z. marina*) showed log linear declines. There was no significant differences between the macroalgal taxa that may be attributed to taxonomy (e.g. red, green & brown algae) and growth form (e.g. filamentous, foliose, fucoid, kelp). This contrasts with the literature, where among macroalgae, brown algae of fucoid and kelp growth forms are expected to be more resistant to degradation ^5,32,55^, having significantly higher proportions of recalcitrant molecules like fucoidans and polyphenolics compared to red and green macroalgae ^32,56,57^. However, a comprehensive meta-analysis ^32^ of known macroalgal degradation rates found mixed evidence of lower degradation rates in brown macroalgae compared to red and green macroalgae, and no clear evidence of polyphenolic content affecting degradation rates. However, it must be noted that none of the studies in the meta-analysis examined degradation in anaerobic sediments. In contrast, the relative overrepresentation of only non-macroalga (*Z. marina*) in this study with time may be due to lignin-rich tracheophyte biomass being more resistant to microbial degradation in anaerobic sediments than macroalgae, which lack lignin altogether ^5^. We caution that the amount of eelgrass biomass input to the sediments used in our study is unknown, and seasonal growth and biomass export of eelgrass in the region is very different from kelp.

In direct contrast to the LBA experiments, in the KB experiment, *S. latissima* COI gene copies increased during the first few weeks of incubation and then plateaued after 4 weeks. The differences in response in eDNA may be due to the different nature of biomass involved and how it was distributed in the sediment. Unlike in the LBA experiment, where fine kelp biomass was well-mixed into the sediment and degraded rapidly, in the KB experiment eDNA may have leaked out from the decaying holdfasts into the surrounding sediment and slowly dispersed by diffusion in pore water ^58^ or possibly meiofaunal-mediated microbioturbation ^59^. The rate of eDNA degradation in the sediment may have then equilibrated with the rate of eDNA release from the decaying holdfasts, as holdfast tissue continued to degrade over the course of the experiment, with a loss of up to 90% of total biomass after 16 weeks. Greater TOC% values in the kelp treatment further support the movement of kelp eDNA into the surrounding sediments, and their subsequent preservation.

In conclusion, we show that both substantial degradation & preservation of sugar kelp eDNA, and by extension, kelp organic carbon, can occur under anaerobic conditions. We also demonstrate strong correlations between multiple eDNA markers from more abundant gene targets. Gene fragment length ratios may have value in characterizing short-term OM degradation in future blue carbon studies. We show that different taxa may exhibit different eDNA degradation dynamics based on metabarcoding data, which highlights the need for complementary quantitative data (e.g. qPCR or dPCR) and context-specific experimental verification (e.g. taxa of interest, sediment type and characteristics). In addition, eDNA findings may differ depending on the form of biomass and the nature of the material being sampled by eDNA. Despite the complex degradation dynamics of macrophyte biomass in anaerobic sediments, we demonstrate that eDNA can be a powerful, taxon-specific, and quantitative tool to complement biogeochemical indicators in blue carbon science.

## Methods

### dPCR assay development and determination of gene copy to biomass relationships

Development of the *S. latissima*-specific ∼150bp mitochondrial (mt) cytochrome oxidase I (COI) digital polymerase chain reaction (dPCR) assay is described in detail in Tan *et al.* (2025) ^17^. Using similar methods, 3 other dPCR assays were developed for a ∼300 bp mt COI fragment, a ∼100 bp chloroplast rbcL fragment, and a ∼120 bp fragment corresponding to a nuclear intergenic region located on Chromosome 7 between regions SJ07277 and SJ07285 of the genome of *S. japonica* ^60^, a close relative of *S. latissima.* With the exception of the 150bp COI assay, where cross-reactivity tests against tissue samples from non-target taxa were conducted, the other three assays were evaluated for cross-reactivity *in-silico*. Primer and probe sequences for all 4 assays can be found in Table S1.

Each assay was run on a series of *S. latissima*-spiked sediment slurries from the spiking experiment described in Tan *et al.*, 2025 ^17^, to examine the relationship between gene copy to added biomass for each marker. Briefly, *S. latissima* biomass was added to 50% sediment slurries in the following amounts by wet mass (0%, 0.01%, 0.05%, 0.1%, 0.5%, 1%, 2% and 5%), in triplicate. eDNA was then extracted from the slurries, without incubation. Given that all 4 assays were designed to have similar optimum annealing temperatures, the cycling conditions were identical for all four assays (Table S1).

### Lyophilized biomass addition (LBA) experiments (LBA: kelp and LBA: mixed)

In mid-April 2023, sediment cores down to approximately 25cm were collected by diver off the Darling Marine Center (DMC) in the Damariscotta River (43.9338, -69.5789) (with a salinity range of 30-32) using acid-washed polycarbonate tubes (10cm internal diameter, 30cm length). Sediment cores were capped underwater with sleeve caps (SC-4 ½, Caplugs) and transported upright to the laboratory to minimize disturbance and penetration of oxygen. Sediment cores were stored in UV-sterilized filtered seawater at 12 °C until use, within 3 days. No visual signs of extensive bioturbation were present in any of the cores.

Prior to sediment core collection, for the LBA: kelp experiment, *S. latissima* blades from healthy individuals were collected off the dock of the DMC. For the LBA: mixed experiment, common beach wrack taxa were collected from the nearby Sand Cove (43.8412, -69.5552). Only fresh, visually healthy specimens of beach wrack drifting in the water column were collected. In addition to *S. latissima*, the following taxa were collected for the LBA: mixed experiment: for angiosperms (i.e. flowering plants), *Zostera marina*; for chlorophytes (i.e. green algae), Ulva sp.; for rhodophytes (i.e. red algae), *Chondrus crispus, Dasysiphonia* sp. and *Rhodomela* sp.; and for ochrophytes (i.e. brown algae), *Fucus vesiculosus* and *Ascophyllum nodosum*. All macrophyte tissue, including *S. latissima*, was lyophilized in a VirTis Advantage XL-70 (SP Scientific) and finely ground with liquid nitrogen in a mortar and pestle, with all equipment cleaned with 1% hydrochloric acid followed by deionized water in between each use. To ensure some degree of genetic variability, the tissue to be lyophilized was sourced from multiple distinct individuals of each taxon. Wet to dry weight ratios were also measured for all taxa, with subsamples dried in a Precision 51221126 drying oven at 70°C (Thermo Fisher Scientific) for 4-5 days.

During the experimental set-up, sediment cores were processed within a portable glove bag (SPILFYTER) flushed with 100% N_2_ gas (MaineOxy) to maintain an anaerobic environment. The upper 5 cm of each sediment core was extruded and discarded, to ensure that only the most hypoxic sediment was used. UV-sterilized filtered seawater from the Damariscotta River (chilled at 12 °C) was bubbled with 100% N_2_ gas until O_2_ levels dropped to approximately zero, as measured with an oxygen microsensor (Unisense). Sediment slurries were then prepared using a 50:50 ratio of sediment and water by volume and either lyophilized *S. latissima* (LBA: kelp) or a mix of the 8 macrophytes (LBA: mixed), homogenized with a clean spatula. No-addition controls were shared between the two experiments, and prepared identically, except for the lack of added macrophyte biomass. Slurries were aliquoted into 50mL plastic conical Falcon tubes (Corning), with triplicates for each week. For the LBA: kelp experiment, each 50mL tube had approximately ∼40mL of slurry and ∼2g of *S. latissima* by wet weight. In the LBA: mixed experiment, each tube instead had ∼0.4g of each macrophyte taxon, by wet weight. Slurries were then transferred to a vinyl anaerobic chamber (Coy Lab products) and incubated anaerobically at 12°C for 16 weeks. For both experiments, all three replicates for at each time point (and controls) were sacrificed on the following weeks after adding lyophilized biomass: 0, 1, 2, 3, 4, 6, 8, 12, and 16, except for week 3, which had no samples for LBA: mixed. Before sampling, tubes were vortexed vigorously with a Vortex Genie 2 (Scientific Industries, Inc) within the anaerobic chamber. Subsamples for eDNA analysis were collected in 5mL conical tubes using 1mL filter pipette tips, with the tips cut off using sterile razor blades. eDNA subsamples were stored at -20 °C prior to eDNA extraction.

### Kelp holdfast burial (KB) experiment

Fresh *S. latissima* holdfasts were acquired from kelp lines at the aquacultural lease of the Darling Marine Center (Maine DMR lease ID: DAM LW2x) in early August 2023 and stored in filtered seawater at 12 °C until use, within 2 days. Holdfasts used for the experiment were shaken dry of excess water, before being measured for wet weight. Holdfast pieces of 4-5g wet weight, and of similar size, shape, remaining stipe length and degree of haptera branching were chosen for consistency. Additional holdfasts were measured for wet weight; and then dried in a drying oven (as described above for lyophilized biomass) for 4-5 days to obtain dry weight.

Additional sediment cores were collected using identical methods to the lyophilized degradation experiment in early August 2023, from the same site. Each sediment core was sub-cored with 3 polycarbonate sediment cores (30cm length x 4cm diameter), which were capped at the bottom with a rubber bung (McMaster-Carr) and secured with electrical tape. Sub-cores were stored in filtered UV-sterilized seawater at 10 °C for the duration of the experiment and aerated individually with a customized manifold made of aquarium tubing, terminating in Luer-lock plastic 18-gauge henna needles.

At the start of the experiment, holdfast pieces were carefully pushed into the sediment in each sub-core, until the haptera of the holdfasts were completely submerged in the sediment, approximately 4cm below the sediment surface. Triplicates were set up to be sacrificed at each time point for the kelp treatment and controls, on the following weeks: 0, 2, 4, 6, 8 and 12.

At each sampling time point, cores were extruded and the upper 4cm section of each core, corresponding to the maximum depth of holdfast burial, was sectioned off and scooped into a weigh boat. The weight of each 4cm section of sediment was measured, including remaining holdfast material. Each sample was then gently broken up, and all visually identifiable holdfast material was removed with tweezers (sterilized in 10% bleach and rinsed with MilliQ water between each use). This holdfast material was carefully rinsed cleaned of sediment with MilliQ water and then weighed for wet weight after blotting off excess water. This holdfast material was then dried and weighed for dry weight (as described above).

Remaining sediment within each slice was homogenized with a sterile scoopula, and then aliquoted into 5mL microfuge tubes, which were stored at -20 °C prior to eDNA extraction. Care was taken to ensure that no *S. latissima* material was visually apparent in the sediment used for eDNA analysis.

### DNA extraction, digital PCR and sequencing

All eDNA subsamples, whether from slurries (LBA: kelp and LBA: mixed) or sediment (KB) were thawed. Prior to DNA extraction, the slurries were homogenized by vortexing and sediments were mixed by hand with a bleach-sterilized scoopula. DNA was extracted from 0.25-0.30 g of wet material using the DNeasy Powersoil Pro Kit (Qiagen) and further cleaned up with the Genomic DNA Clean & Concentrator Kit (Zymo Research), following manufacturers’ protocols. Extracted DNA was initially quantified with a Nanodrop ND-1000 (Thermo Fisher Scientific). DNA was then diluted to 5-10 ng/uL and re-quantified using a Qubit 3.0 Fluorometer and the dsDNA BR kit (ThermoFisher Scientific) before downstream use.

The 4 digital PCR (dPCR) assays were run on samples from the kelp and control samples of the LBA experiment. Only the 150bp COI assay was run on the mixed macrophyte samples from the LBA experiment, and the samples from the KB experiment. dPCR assays were conducted on the QIAcuity Four dPCR system (Qiagen), using 8.5k-partition Qiacuity Nanoplates (Table S1).

For metabarcoding, DNA extracts were sent to the University of New Hampshire’s Hubbard Center for Genome Studies (UNH HCGC) for amplicon sequencing of the 16S V4-V5 (primer set: 515/926R ^61^) and 18S V9 (primer set: 1391F/1510R ^62,63^) regions on the Novaseq 6000 platform (Illumina), following Earth Microbiome protocols ^64^. Samples were sequenced to a read depth of about 200k reads per sample.

### TOC% and TN

Frozen sediment from the LBA: kelp and KB experiments was thawed, aliquoted into weigh boats and measured for wet weight, before being dried in an incubator (Model 12-140 Incubator, Quincy Lab Inc.) at 70 deg C for ∼5 days, after which dry weight was measured. Samples were then carefully ground with a mortar and pestle, cleaned with 3.7% hydrochloric acid (HCl) and dried between each sample. Ground samples were aliquoted into glass vials and carefully acidified by dripping 1M HCl, until no remaining effervescence was noted. Acidified samples, including samples for TN, were rinsed with deionized water, and once again dried and ground as described above. Samples were weighed with a Supermicro S4 microbalance (Sartorius), packaged into tin capsules and sent to the Robinson Lab of the University of Rhode Island Graduate School of Oceanography for Elemental Analysis-Isotope Ratio Mass Spectrometry (EA-IRMS) on a Delta V-Advantage Isotope Ratio Mass Spectrometer (Thermo Scientific) coupled to a ECS 4010 Elemental Analyzer (Costech Analytical Technologies Inc.).

### Data analysis

Except for initial sequence read processing, all data analysis was conducted in R (version 4.4.2) ^65^. All underlying code, data, and parameters used for the analyses and plots in this study, as well as the output of any statistical tests, can be found on Zenodo at DOI: <u>10.5281/zenodo.18840678</u>. All raw 16S and 18S reads used in this study have been deposited with the National Center for Biotechnology Information (NCBI) under BioProject ID <u>PRJNA1430818</u>. For each of the four dPCR assays, linear regressions were conducted on log gene copy number against added *S. latissima* biomass in the spiking experiment samples, in terms of % added dry *S. latissima* carbon. Wilcoxon signed rank tests were used to examine if there were significant differences between gene copy numbers, for pairwise comparisons between each of the assays.

For the metabarcoding data, 16S and 18S reads were processed using a custom dada2 ^66^ pipeline on the Bigelow Laboratory for Ocean Sciences (BLoS)’s supercomputing cluster. The following analyses were primarily conducted using the phyloseq ^67^ and vegan ^68^ packages, with all plots made with ggplot2 ^69^. For the 16S dataset and 18S dataset, all samples were rarefied to 15 000 reads and 990 reads, respectively. For the 16S dataset, amplicon sequence variants (ASVs) corresponding to chloroplasts were agglomerated at the order level and calculated for every sample. ASVs corresponding to the added 8 macrophyte taxa (from the classes Phaeophyceae, Florideophyceae, Embryophyceae, and Ulvophyceae) were subset from the total 18S dataset and used for downstream analyses. The top 10 ASVs from this subset of macroalgal classes were found to correspond to the added macrophyte taxa, as verified by nucleotide BLAST against the NCBI nr/nt database ^70^. While each of the added macrophyte taxa was predominantly represented by a single ASV (>95% relative abundance), ASVs were agglomerated at the genus level to account for less abundant ASV variants from the same species. Similarly to the chloroplast reads, read abundance for each taxon was then calculated for each sample.

Kruskal-Wallis (KW) tests were conducted to test for significant differences in the eDNA and nutrient metrics based on treatment type, followed by Dunn’s tests. Benjamin-Hochberg correction was applied for KW tests comparing three treatment levels (i.e. kelp, mixed, control in the LBA experiment).

To examine the correlations between the different quantitative (150bp COI, 300bp COI, rbcL, NC) and qualitative (16S chloroplast reads and 18S *S. latissima* reads) eDNA variables, and non-eDNA markers (TOC% and TN%), correlation plots were made with the Hmisc ^71^ and corrplot ^72^ R packages, solely on data from the kelp treatment of the LBA experiment. When examining TOC% and TN% values in the kelp treatment, they were subtracted from the mean value in the controls and referred to as added TOC% and TN% respectively.

To quantify changes with time of different metrics, data from the kelp treatment (for both LBA and KB experiments) was subset, and various models were fit using the nls function in the base R stats package. Linear or non-linear least squares (NLS) models were chosen for each measured variable depending on best fit; based on analysis of variance (ANOVA) comparisons with each other and a null model of gradient = 1 (representing a linear fit). For the NLS models, various common ecological relevant models approximating broad patterns seen in the eDNA and non-eDNA data were evaluated using ANOVA as described above [declines to an asymptote (e.g. exponential decline, power-law exponential decline, negative Gompertz); increases to an asymptote (e.g. logistic, Michaelis-Menten, Gompertz)], but in the end, final models were chosen based on a mix of ANOVA results and parsimony for the sake of interpretability.

## Supporting information

Supplementary Documents (all)

Figure 3 (higher-res)

Figure 2 (higher-res)

Figure 1 (higher-res)

## Acknowledgements

We would like to acknowledge the following: from the Darling Marine Center, Sean O’Neill for assisting with sediment core collection, Margaret Estapa for providing manuscript feedback, and Adam St. Gelais for contributing kelp holdfasts; and from the Bigelow Laboratory for Ocean Sciences, Amy Doiron for assisting with DNA extractions, and Peter Countway for use of laboratory facilities and associated support.

