## Supplementary Documents (all) for "Assessing the degradation dynamics of sugar kelp in anaerobic marine sediment using environmental DNA"

Table S1: Primer and probe sequences and cycling conditions for the 4 dPCR assays in this paper: 150bp COI (Tan *et al.*, 2025 ^141^), 300bp COI, rbcL and NC. All probes are TaqMan -minor groove binder (MGB) probes, listed without dyes. As mentioned in Tan *et al.* (2025), master mix composition is as follows: 3.4 µL of Qiacuity Probe Kit Master Mix (Qiagen), 0.544 µL of the probe, 1.088 µL each of both forward and reverse primers, 5.88 µL of nuclease-free water, and 4 µL of sample, for a total reaction volume of 16 µL. dPCR thermal cycler conditions were as follows: initial denaturation at 95 ℃, for 2 minutes, followed by 40 cycles: denaturation at 95 ℃ for 15 seconds, and combined annealing and extension at 60 ℃ for 1 minute.

| Assay | Forward primer (F) | Reverse primer (R) | Probe (P) | Approx. amplicon length (bp) |
| --- | --- | --- | --- | --- |
| 150bp COI | CCC CTC TTT AAT CTT GCT TC | CCT GAA AGA TGG AGA CTA AAT ATA G | AGC GTC CTC ATT GGA | 150 |
| 300bp COI | CCC TCT TTA ATC TTG CTT CT | AAT TCC TAT CGG TTAA TAA CAT TG | CGT CCT CAT TGG TAG AAT | 300 |
| rbcL | GGA CTT CTA AAT TAT TCC CTC | TCG AGT TCT TTC CTG TAC | AGA CAT ATA GAT GCT CCG AAG CCA GC | 100 |
| NC | ACC ACC AGT AGA ATT TGA G | CAT GCT TGT TCC TTG AGA | AAC GCA CCA CC GGC TGC AAA | 120 |

Table S2: Linear regression results for the 4 dPCR assays in the study. For the equations, y = log gene copy number per gram of dry sediment, x = % added dry *S. latissima* carbon. Bracketed values in the equations represent standard error for the slope term.

| Assay | Source | Equation | R^2^ | p-value |
| --- | --- | --- | --- | --- |
| 150bp COI* | Mitochondrial | y = 1.33*10^10^ (+/- 6.23* 10^8^) * x | 0.93 | <1e^-10^ |
| 300bp COI | Mitochondrial | y = 8.35 * 10^9^ (+/- 6.23*10^8^) * x | 0.94 | <1e^-10^ |
| rbcL | Chloroplast | y = 2.69 *10^10^ (+/- 1.19*10^9^) * x | 0.93 | <1e^-10^ |
| NC | Nuclear | y = 2.17 * 10^8^ (+/- 8.01*10^6^) * x | 0.95 | <1e^-10^ |

*Reported in Tan *et al.,* 2025

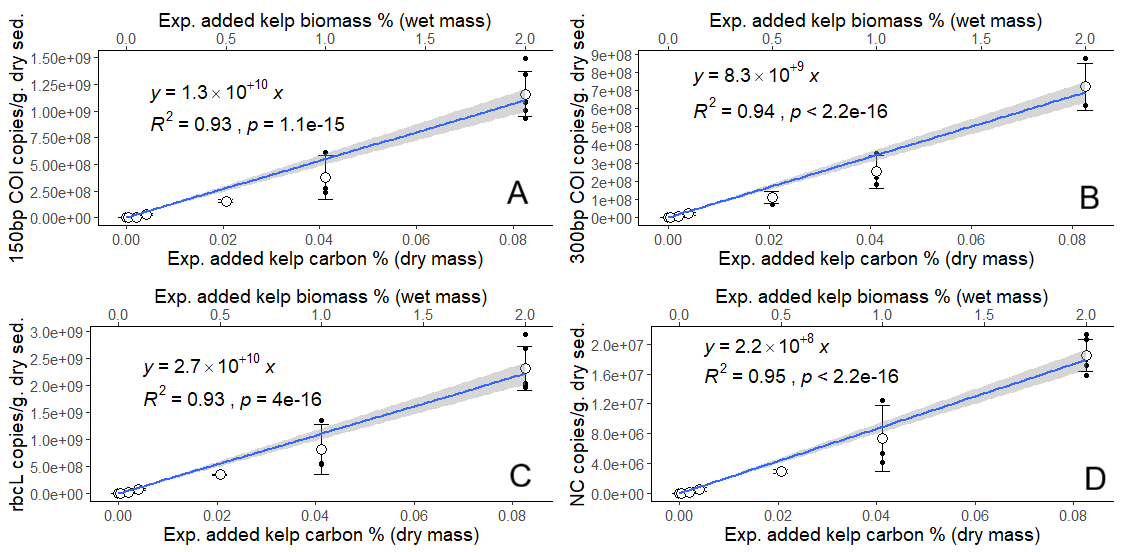

Figure S1: Linear regressions of gene copy number against experimentally added *S. latissima* biomass in the kelp spiking experiment, for 150bp COI (Tan *et al.,* 2025), 300bp COI, rbcL and NC. The y-axis refers to gene copy numbers normalized per gram of dried sediment slurry, and the primary x-axis (lower) refers to percentage added dry *S. latissima* carbon per gram of dried sediment slurry. The secondary x-axis (upper) refers to percentage added wet *S. latissima* biomass per gram of wet sediment slurry. Regression equations, R^2^ values and p-values are provided on each plot.

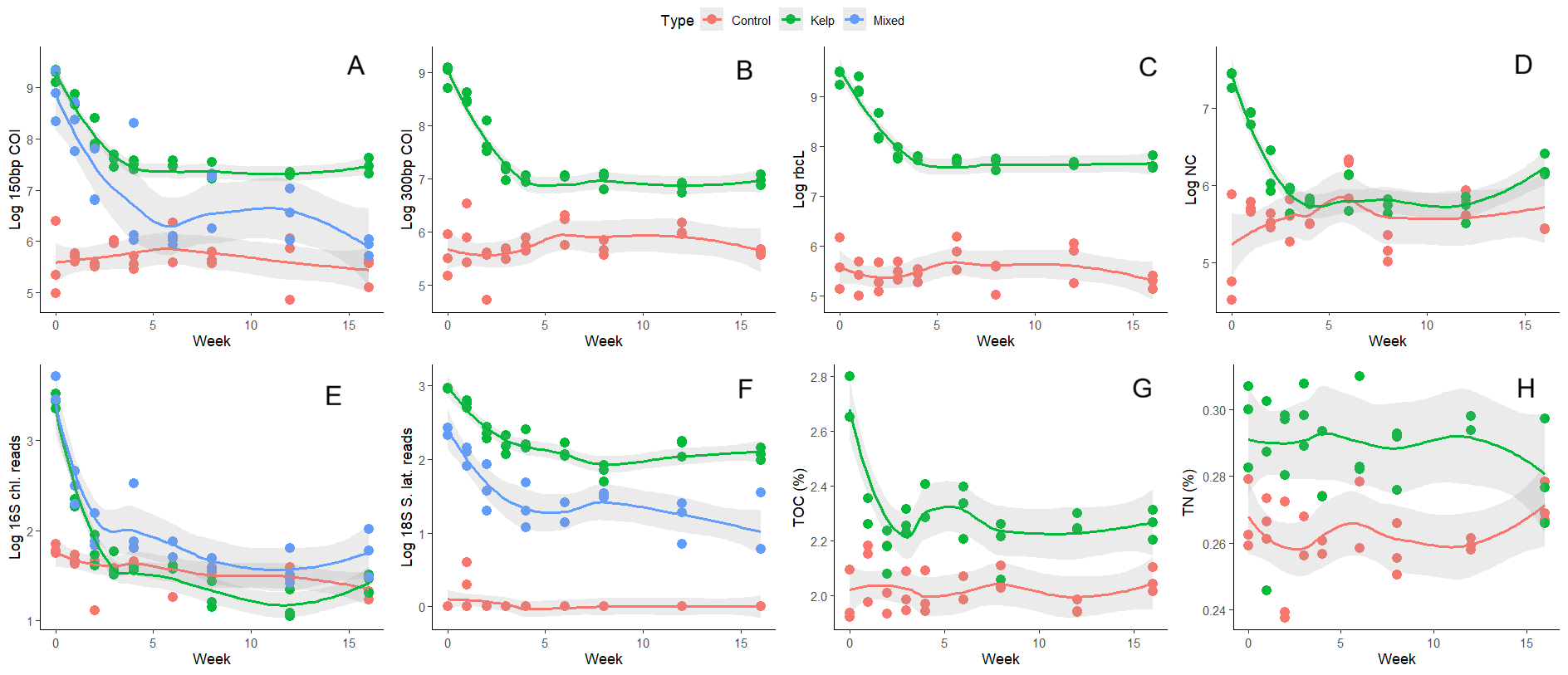

Figure S2: Quantitative and qualitative eDNA metrics, as well as TOC% and TN% for LBA: kelp (Kelp) and LBA: mixed experiments (Mixed), and the controls. Smoothed means have been drawn through the points for each treatment type for each metric, with the surrounding gray outline representing the 95% confidence interval.

Table S3: Results of Kruskal-Wallis tests and post-hoc Dunn’s tests done to examine significant differences between treatments in the combined LBA: kelp + LBA: mixed dataset. P-values for all Kruskal-Wallis tests are annotated after the X^2^ value (p < 0.0001 = ***). Mean values for each treatment are given in log values for the eDNA metrics.

| Metric | Kruskal-Wallis X^2^ | df | Dunn’s test results | Treatment mean +/- SD |
| --- | --- | --- | --- | --- |
| Log 150bp COI ^#^ | 50.31 *** | 2 | Kelp > Mixed > Control | K: 7.858 +/- 0.671  M: 7.057 +/- 1.117  C: 5.670 +/- 0.361 |
| Log 300bp COI | 39.7636 *** | 1 | Kelp > Control | K: 7.461 +/- 0.749  C: 5.733 +/- 0.357 |
| Log rbcL | 39.7636 *** | 1 | Kelp > Control | K: 8.133 +/- 0.679  C: 5.507 +/- 0.331 |
| Log NC | 16.458 *** | 1 | Kelp > Control | K: 6.183 +/- 0.575  C: 5.578 +/- 0.413 |
| Log 16S chloroplast reads ^#^ | 13.5009 *** | 2 | Kelp ~ Mixed^†^ > Control | K: 1.778 +/- 0.696  M: 2.056 +/- 0.600  C: 1.568 +/- 0.182 |
| Log 18S *S. latissima* reads ^#^ | 62.463 *** | 2 | Kelp > Mixed > Control | K: 2.300 +/- 0.348  M: 1.549 =/- 0.502  C: 0.0446 +/- 0.137 |
| TOC% | 33.2436 *** | 1 | Kelp > Control | K: 2.271 +/- 0.244  C: 2.010 +/- 0.115 |
| TN% (acidified) | 28.0355 *** | 1 | Kelp > Control | K: 0.285 +/- 0.0284  C: 0.271 +/- 0.0441 |

^#^Also examined in the Mixed treatment.

^†^There was no statistically significant difference between the treatment pairs.

Table S4: Summary of models fit to the eDNA metrics, added TOC% and added TN% (i.e. subtracting mean values from the controls), as shown for the LBA: kelp experiment in Figure 1. Model significance is annotated in the log likelihood value column (p < 0.0001 = ***). Estimated T_50_ values (i.e. time taken to decay to 50% of the initial value) are calculated based on the non-log transformed starting values at week 0, using the equations in the table (the 150bp:300bp COI ratio has been omitted since it increases with time).

| Marker (Y) | Model Type | Equation | Log Likelihood | Residual Standard Error | Total mean % change (W0 to W16) | Est. T_50_ (weeks) |
| --- | --- | --- | --- | --- | --- | --- |
| Log 150bp COI | Exponential | Y ~ 1.984 * (exp(-0.526*Week)) + 7.345 | 9.911 *** | 0.178; 24 df | -98.3% | 0.415 |
| Log 300bp COI | Exponential | Y ~ 2.195 *(exp(-0.499*Week)) + 6.874 | 5.068 *** | 0.213; 24 df | -98.9% | 0.438 |
| Log rbcL | Exponential | Y ~ 1.969 * (exp(-0.477*Week)) + 7.591 | 4.918 *** | 0.214, 24 df | -98.2% | 0.559 |
| Log NC | Exponential | Y ~ 1.614 * (exp(-0.734*Week)) + 5.845 | 0.270 *** | 0.254; 24 df | -92.8% | 0.366 |
| 150bp: 300bp COI | Logistic | Y ~ 3.082/ (1 + 0.780*exp(-0.471*Week)) | -11.584 *** | 0.402; 23 df | +53.0% | NA |
| Log 16S chloroplast | Exponential | Y ~ 2.067* (exp(-0.732*Week)) + 1.349 | 12.290 *** | 0.158; 22 df | -99.0% | 0.202 |
| Log 18S *S. latissima* | Exponential | Y ~ 0.976* (exp(-0.478*Week)) + 2.032 | 16.024 *** | 0.142; 24df | -87.2% | 0.904 |
| Added TOC% | Exponential | Y ~ 0.497* (exp(-2.611*Week)) + 0.243 | 27.440 *** | 0.0861; 22 df | -66.2% | 0.358 |
| Added TN% | Linear | Y ~ -0.000371*Week + 0.0233 | 76.802 | 0.0103; 22 df | -37.7% | 28.3 |

Table S5: Spearman’s correlation analysis results for the eDNA metrics and added TOC% and added TN% in the *S. latissima*-only LBA: kelp experiment. Values are rho values, reflecting the strength of correlations (from -1 to +1). All pairwise combinations that were statistically significant have been marked out (*: p < 0.05; **: p < 0.01; ***: p<0.001).

|  | 300bp mt. COI | chl. rbcL | NC | Added TOC% | Added TN% | 16S chl. | 18S *S. lat.* |
| --- | --- | --- | --- | --- | --- | --- | --- |
| 150bp mt. COI | 0.95 *** | 0.95 *** | 0.73 *** | 0.35 | 0.25 | 0.85 *** | 0.84  *** |
| 300bp mt. COI |  | 0.94 *** | 0.73 *** | 0.29 | 0.19 | 0.80 *** | 0.75 *** |
| chl. rbcL |  |  | 0.62 *** | 0.37 | 0.24 | 0.83 *** | 0.85 *** |
| NC |  |  |  | 0.30 | 0.05 | 0.58 ** | 0.56 ** |
| TOC (%) |  |  |  |  | 0.28 | 0.44 * | 0.36 |
| TN (%) |  |  |  |  |  | 0.08 | 0.24 |
| 16S chl. |  |  |  |  |  |  | 0.81 *** |

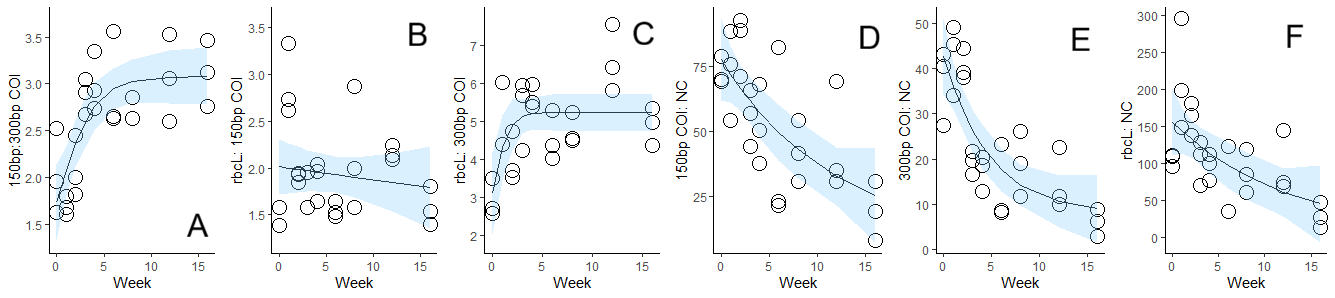

Figure S3: Plots of models fit to the gene ratios in the LBA: kelp experiment, with 95% confidence intervals in light blue.

Table S6: Summary of models fit to the gene ratios analyzed in the LBA: kelp experiment, in Figure S3. Note that the 150bp:300bp COI row is identical to that in Table S3. Model significance is annotated in the log likelihood value column (p < 0.01 = **; p < 0.0001 = ***)

| Gene ratios (Y) | Model Type | Equation | Log Likelihood | Residual Standard Error | Total mean % change (W0 to W16) |
| --- | --- | --- | --- | --- | --- |
| 150bp:300bp COI | Logistic | Y ~ 3.082/ (1 + 0.780*exp(-0.471*Week)) | -11.584 *** | 0.402; 23 df | +53.0% |
| rbcL: 150bp COI | Linear | Y ~ -0.014*Week + 2.011 | -17.951 | 0.489; 25 df | +9.1% |
| rbcL:300bp COI | Logistic | Y ~ 5.226/ (1 + 0.691*exp(-1.166*Week)) | -34.564 ** | 0.923, 24 df | +67.3% |
| 150bp COI: NC | Exponential | Y ~ 70.514 * (exp(-0.086*Week)) + 7.370 | -114.119 ** | 17.630; 24 df | -72.0% |
| 300bp COI: NC | Exponential | Y ~ 34.895* (exp(-0.213*Week)) + 7.877 | -94.039 *** | 8.355; 24 df | -73.5% |
| rbcL: NC | Exponential | Y ~ 151.686* (exp(-0.081*Week)) + 4.184 | -142.417 *** | 50.130; 24 df | -83.9% |

Table S7: Summary of models fit to the log 18S read counts for each of the added macrophyte taxa in the LBA: mixed experiment. Total mean % change from start to end of the experiment (W0-W16) is with reference to the non-log values of the relevant metrics. Model significance is annotated in the log likelihood value column (p < 0.05 = *; p < 0.01 = **; p < 0.0001 = ***). Estimated T_50_ values are calculated based on the non-log transformed starting values at week 0, using the equations in the table (*Z. marina* and *Ulva* sp. have been omitted since the models are non-significant, and estimated T_50_ values are negative).

| Taxon [log 18S reads] (Y) | Model Type | Equation | Log Likelihood | Residual Standard Error | Total mean % change (W0 to W16) | Est. T_50_ (weeks) |
| --- | --- | --- | --- | --- | --- | --- |
| *Saccharina latissima* | Exponential | Y ~ 1.223* (exp(-0.437*Week)) + 1.185 | 0.165 *** | 0.258; 20 df | -93.7% | 0.676 |
| *Fucus vesiculosus* | Linear | Y ~ -0.0379*Week + 2.136 | -3.807 ** | 0.294; 23 df | -60.2% | 9.474 |
| *Ascophyllum nodosum* | Linear | Y ~ -0.0634*Week + 2.250 | -4.328 *** | 0.313; 19 df | -83.9% | 7.329 |
| *Zostera marina* | Linear | Y ~ 0.0112*Week + 1.542 | -0.916 | 0.263, 22 df | +386.9% | NA |
| Family: *Dasyaceae* | Exponential | Y ~ 1.0401* (exp(-0.515*Week)) + 1.248 | -4.677 *** | 0.318; 20 df | -88.2% | 0.654 |
| Genus: *Rhodomela* | Linear | Y ~ -0.0260*Week + 1.0421 | -2.347 * | 0.285; 19 df | -73.5% | 11.107 |
| *Chondrus crispus* | Linear | Y ~ -0.0812*Week + 1.876 | -5.367 *** | 0.320; 21 df | -98.2% | 1.736 |
| Genus: *Ulva* | Linear | Y ~ -0.0194*Week + 1.0492 | -4.854 | 0.3251; 18 df | -73.5% | NA |

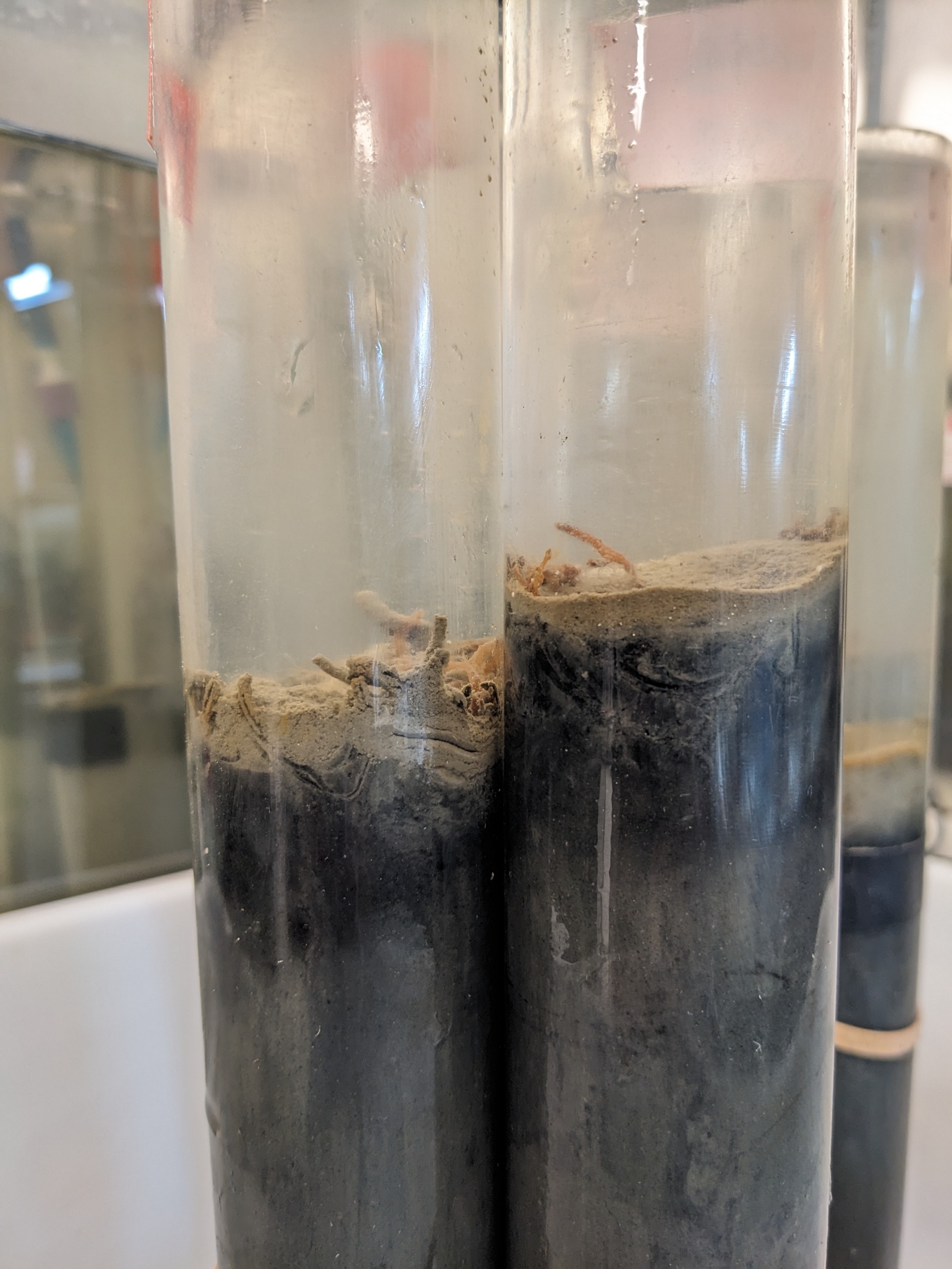

Image S1: The experimental set-up in the KB experiment after 16 weeks of incubation. Sediment at depths where the holdfasts were buried often turned dark with time, indicating sulfate reduction and subsequent formation of iron sulfides. In some cases, the sediment surface developed white patches indicative of sulfur-oxidizing bacteria, but they did not persist for more than a few weeks. Oxygen measurements made in the water column above the cores found fully oxic conditions throughout the experiment.

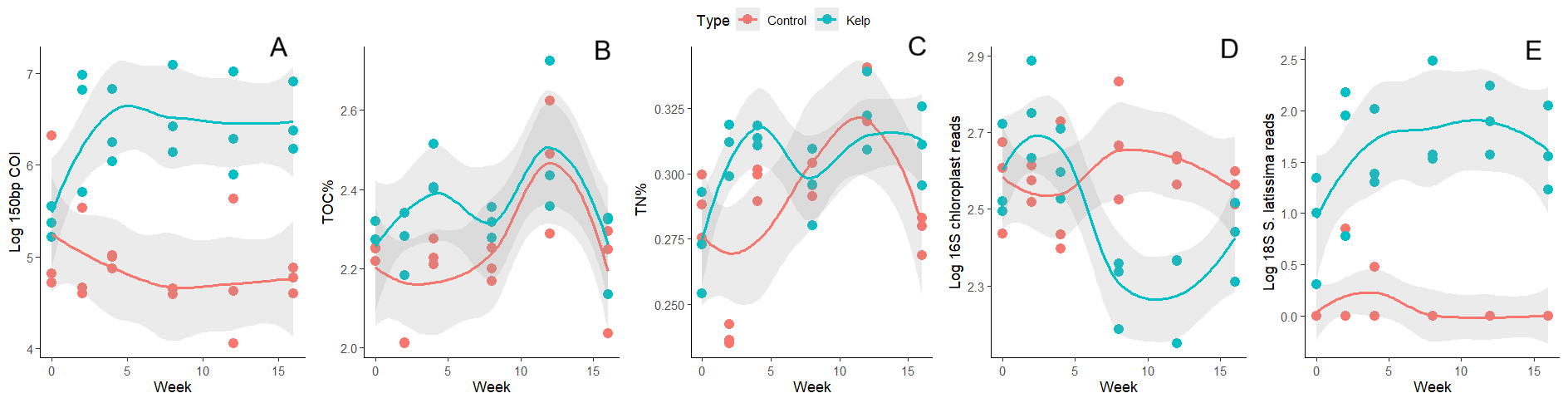

Figure S4: Plots of eDNA metrics, TOC% and TN% in the KB experiment, with smoothed means and associated 95% confidence intervals.

Table S8: Results from the KB experiment of Kruskal-Wallis tests and post-hoc Dunn’s tests to examine significant differences between treatments. P-values for all Kruskal-Wallis tests are annotated after the X^2^ value (p < 0.05 = *; p < 0.01 = **; p < 0.0001 = ***). Mean values for each treatment are given in log values for the eDNA metrics.

| Metric | Kruskal-Wallis X^2^ | df | Dunn’s test results | Treatment mean +/- SD |
| --- | --- | --- | --- | --- |
| Log 150bp COI | 21.631 *** | 1 | Kelp > Control | K: 6.280 +/- 0.582  C: 4.889 +/- 0.502 |
| TOC% | 8.2716 ** | 1 | Kelp > Control | K: 2.351 +/- 0.131  C: 2.238 +/- 0.152 |
| TN% | 3.8478 * | 1 | Kelp > Control | K: 0.304+/- 0.0204  C: 0.288 +/- 0.0300 |
| Log 16S chloroplast reads | 2.7067 | 1 | NA | K: 2.493 +/- 0.199  C: 2.584 +/- 0.108 |
| Log 18S *S. latissima* reads | 27.737 *** | 1 | Kelp > Control | K: 1.578 +/- 0.552  C: 0.0735 +/- 0.223 |

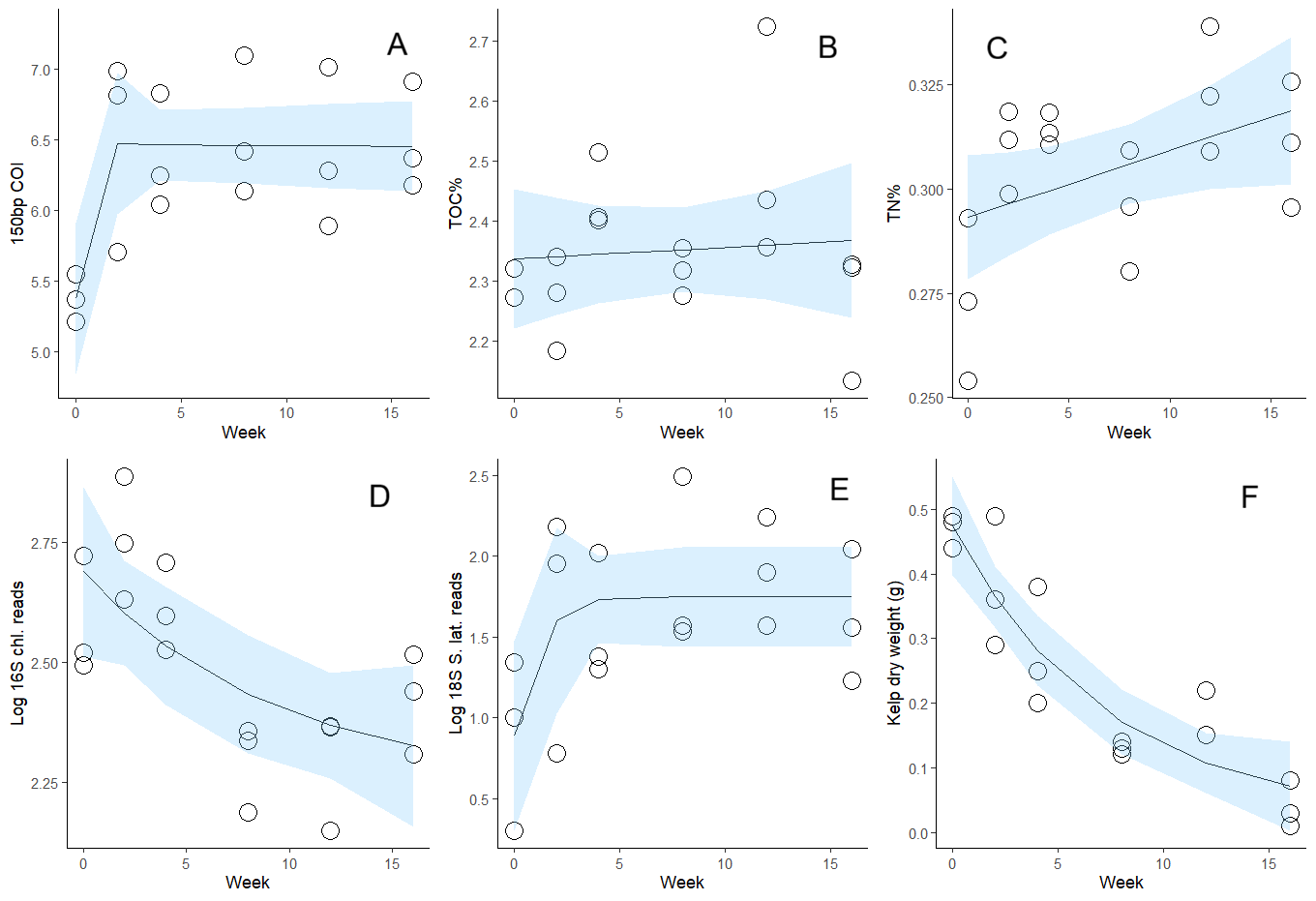

Figure S5: Models fit to the various eDNA and nutrient metrics, and *S. latissima* dry mass in the kelp treatment of the KB experiment, over time, with 95% confidence intervals in light blue.

Table S9: Summary of models fit to the eDNA metrics, TOC% and TN%, and *S. latissima* dry mass in the KB experiment. Total mean % change from start to end of the experiment (W0-W16) is with reference to the non-log values of the relevant metrics. Model significance is annotated in the log likelihood value column (p < 0.05 = *; p < 0.01 = **; p < 0.0001 = ***). Estimated T_50_ values have been calculated for (non-log transformed) 16S chloroplast read counts and *S. latissima* dry weight using the equations in the table, as they are the only variables which declined with time.

| Marker (Y) | Model Type | Equation | Log Likelihood | Residual Standard Error | Total mean % change (W0 to W16) |
| --- | --- | --- | --- | --- | --- |
| Log 150bp COI | Michaelis-Menten | Y ~ 5.377 + ((1.076*Week)/(-0.034 + Week)) | -8.901 ** | 0.435; 15 df | +1494.4% |
| TOC% | Linear | Y ~ 0.0002*Week + 2.337 | 10.990 | 0.135; 15 df | +4.2% |
| TN% (acid.) | Linear | Y ~ 0.0002*Week + 0.293 | 47.160 | 0.0187; 16 df | +13.7% |
| Log 16S chloroplast | Exponential | Y ~ 0.442* (exp(-0.108*Week)) + 2.248 | 9.263 * | 0.158; 15 df | -31.1% (T_50_: 21.623 weeks) |
| Log 18S *S. latissima* | Logistic | Y ~ 1.747/(1 + 0.970*exp(-1.182*Week)) | -10.566 * | 0.477; 15 df | +382.4% |
| *S. latissima* dry weight (g) | Exponential | Y ~ 0.452* (exp(-0.139*Week)) + 0.022 | 24.786 *** | 0.0669; 15 df | -91.5% (T_50_: 5.413 weeks) |

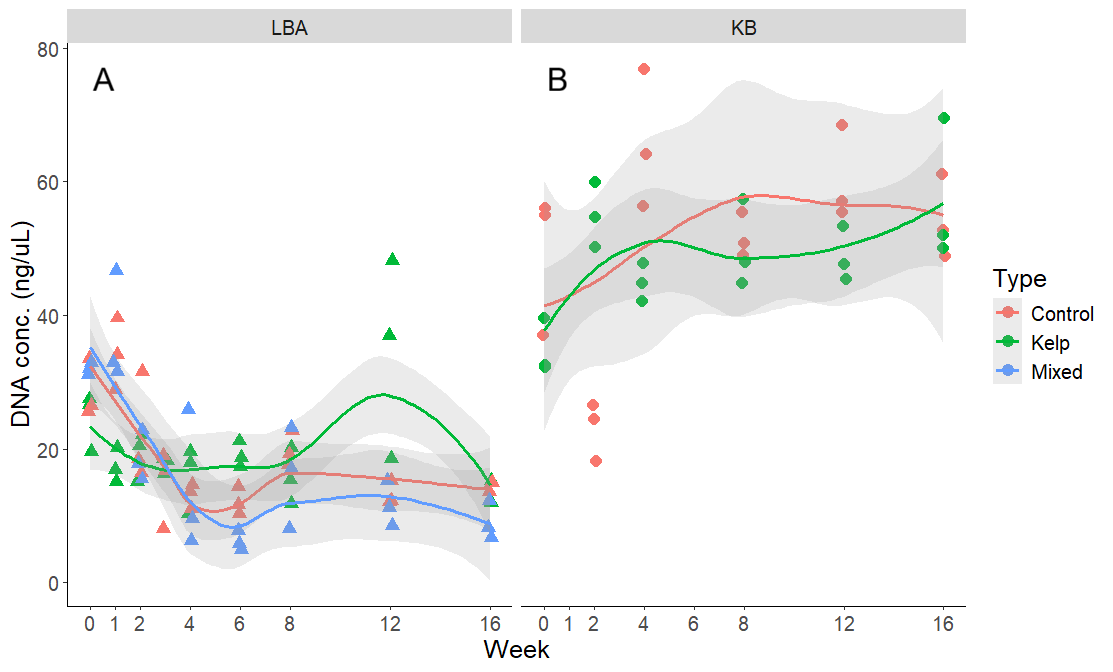

Figure S6: Concentration of extracted DNA from samples from the LBA (LBA: kelp and LBA: mixed combined) and KB experiments. Smoothed means are drawn for each treatment type, with 95% confidence intervals in gray. Kruskal-Wallis tests found treatment type to be non-significant for both the LBA (X^2^ = 2.0337, df = 2, p = 0.3617) and KB experiments (X^2^ = 1.9379, df = 1, p = 0.1639).
