## Supplementary figures and images for "Assessing the degradation dynamics of sugar kelp in anaerobic marine sediment using environmental DNA"

### Figure 1 (higher-res)

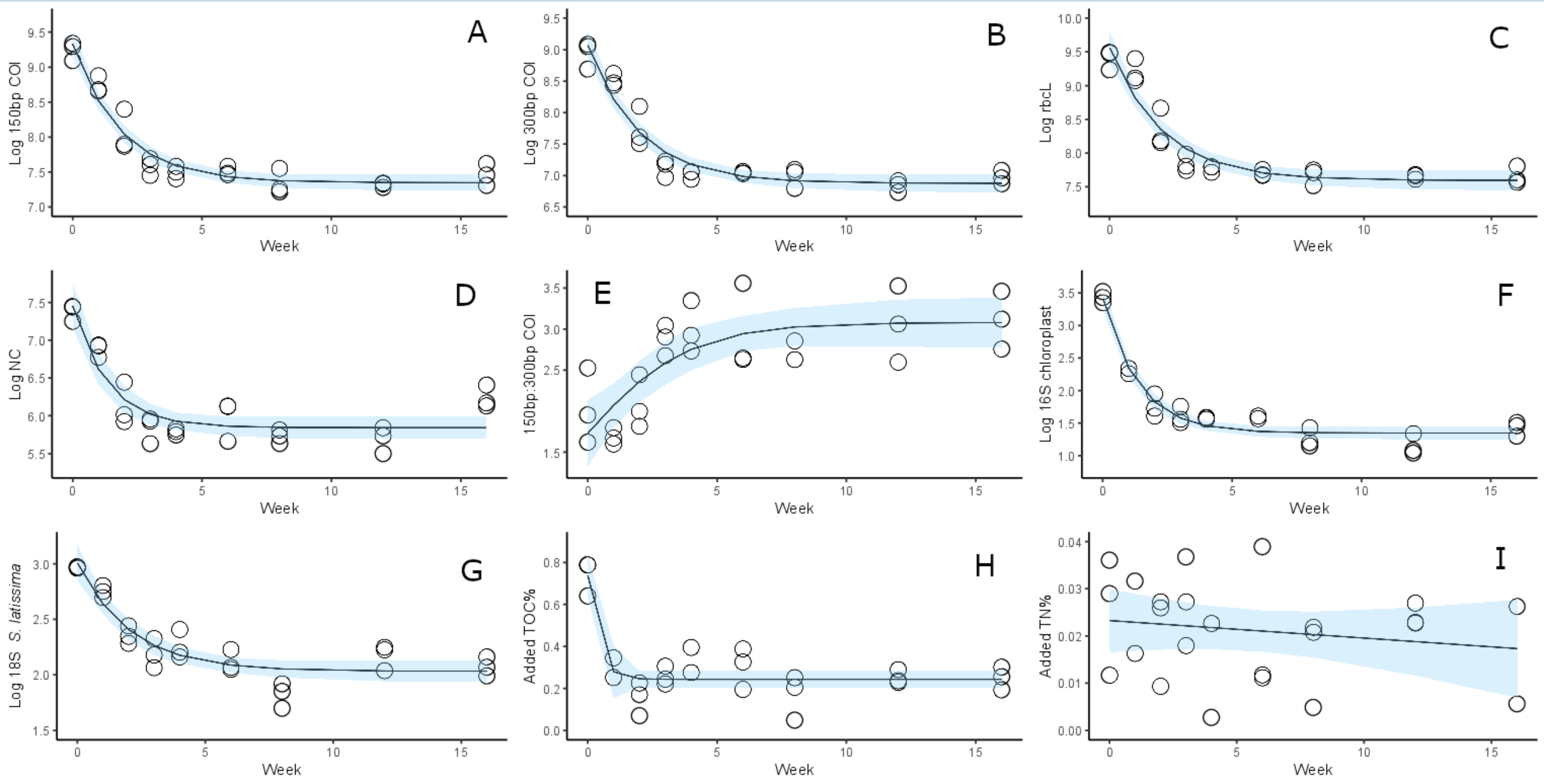

### Figure 2 (higher-res)

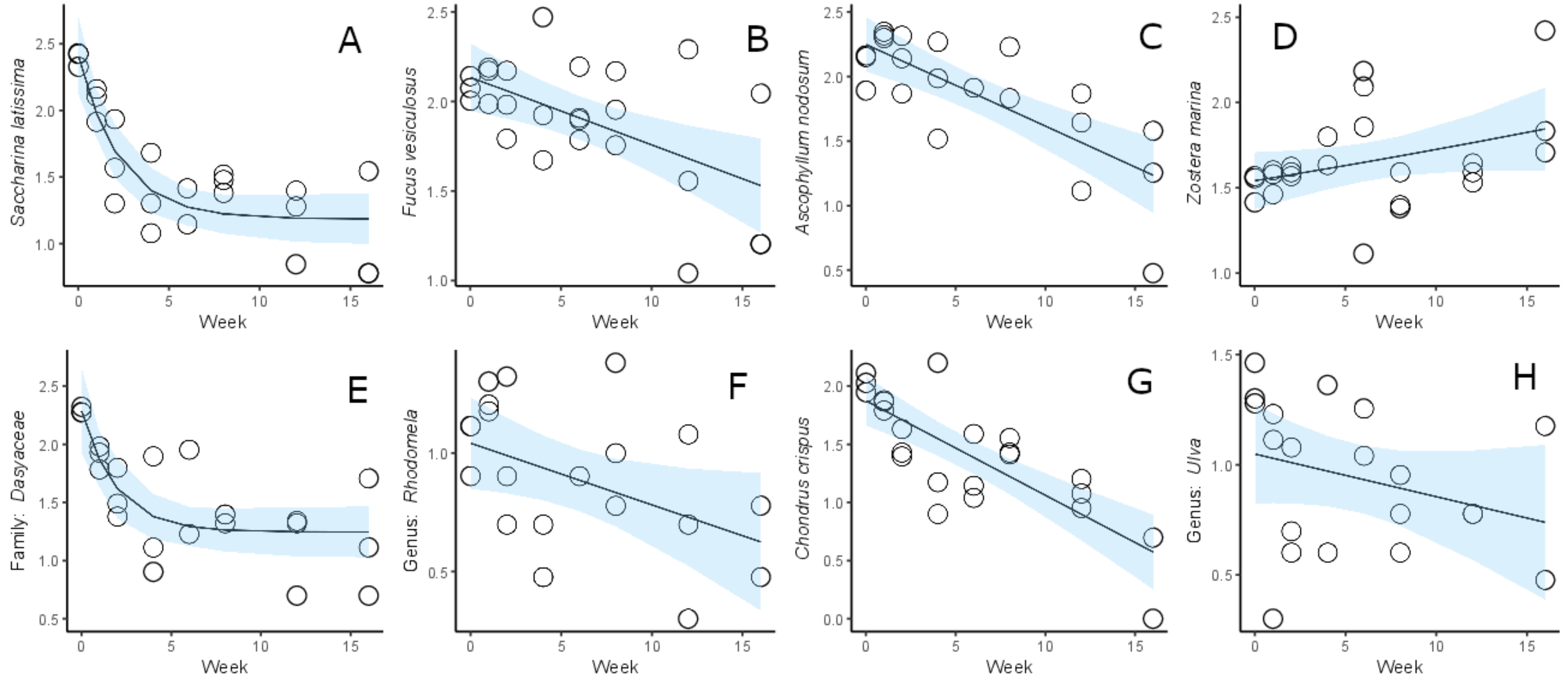

### Figure 3 (higher-res)

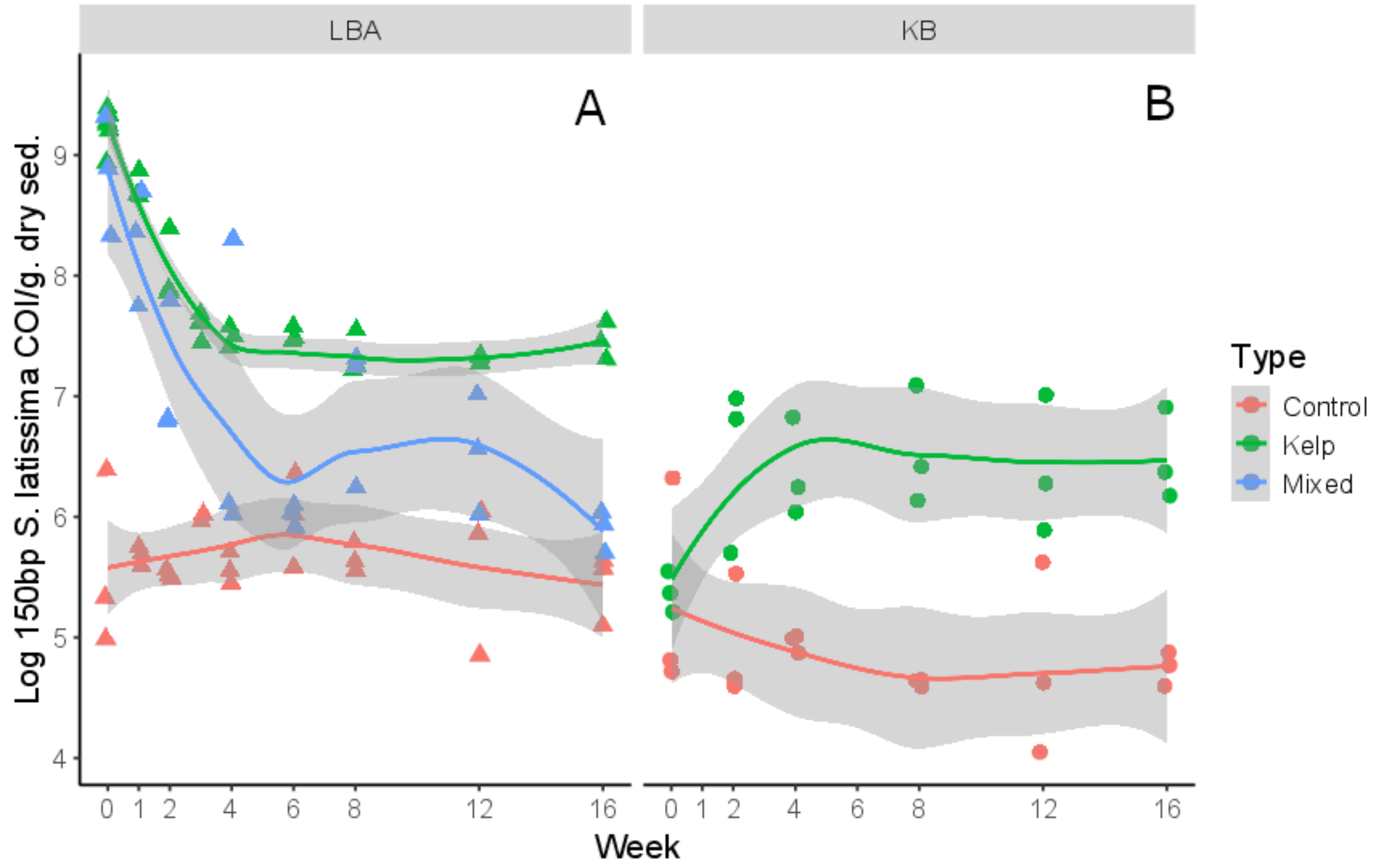
